# Light prevents pathogen-induced aqueous microenvironments via potentiation of salicylic acid signaling

**DOI:** 10.1101/2022.08.10.503390

**Authors:** Gaële Lajeunesse, Charles Roussin-Léveillée, Sophie Boutin, Élodie Fortin, Isabelle Laforest-Lapointe, Peter Moffett

**Affiliations:** Centre SÈVE, Département de Biologie, Université de Sherbrooke, Sherbrooke, Québec, Canada

## Abstract

Upon establishment of an infection, many plant pathogens induce an aqueous microenvironment in the extracellular space of their host, resulting in water-soaked lesions. In the case of *Pseudomonas syringae* (*Pst*), this is accomplished through the activity of water-soaking effectors that stimulate abscisic acid (ABA) production and signaling, which results in stomatal closure. This reduces transpiration and induces a microenvironment favorable for bacterial growth. Stomata are also highly sensitive to environmental conditions, including light and circadian rhythm. Here, we show that a period of darkness is required for water-soaking, and that a constant light regime abrogates the water-soaking activity of *Pst* effectors. Additionally, we show that constant light induces resistance against *Pst* and that this effect requires salicylic acid (SA). An increase in SA production upon infection under constant light did not affect effector-induced ABA signaling, but rather abrogated ABA’s ability to induce stomatal closure. Indeed, under normal diurnal light regimes, application of a SA analog is sufficient to prevent the ability of the pathogen to induce stomatal closure and a water-rich niche in the apoplast. Our results provide a novel approach to interfering with a common virulence strategy, as well as providing a physiological mechanism by which SA functions in defense against certain pathogens.

## Introduction

The outcome of a plant-pathogen interaction is determined by multiple factors. These include environmental conditions, as well as plant defense mechanisms, and the ability of pathogens to avoid or overcome the latter (Cheng et al., 2019; Nishad et al., 2020). Plants encode a large variety of pattern recognition receptors that can detect microbe-associated molecular patterns (MAMPs) and induce pattern-triggered immunity (PTI) (Han, 2019; Nishad et al., 2020; Yuan et al., 2021). Early PTI responses include acidification of the apoplast, production of reactive oxygen species (ROS) and transcriptional reprogramming (Han, 2019; Nishad et al., 2020; Yuan et al., 2021). Later responses include biosynthesis of defense-related phytohormones such as salicylic acid (SA), jasmonic acid (JA) and ethylene (Nishad et al., 2020; Yuan et al., 2021). In turn, a key virulence mechanism used by pathogens is the translocation of effector proteins into host plant cells, many of which have been shown to inhibit PTI (Toruño et al., 2016). Plant pathogenic bacteria possess a needle-like structure, commonly referred to as the type-3 secretion system (T3SS), that delivers effector proteins which promote disease progression in their hosts (Collmer et al., 2000; Lee et al., 2012). However, inhibition of defense responses is not sufficient for optimal pathogen proliferation and additional effector activities are required for the establishment of favorable microenvironments in the apoplast (Gentzel et al., 2022; Hu et al., 2022b; Roussin-Léveillée et al., 2022; Xin et al., 2016, 2018).

Plant stomata are often targeted by pathogenic effector proteins or toxins as these structures represent natural breaches for pathogen entry (Melotto et al., 2006; Xin et al., 2018). Indeed, an important aspect of early response to pathogens involves stomatal immunity, wherein PTI responses result in stomatal closure (Melotto et al., 2006, 2017). To counteract stomatal immunity, *Pseudomonas syringae* (*Pst*) produces a JA-mimicking phytotoxin called coronatine (COR). COR has been reported to induce stomatal reopening to allow the pathogen to gain access to the apoplastic space (Panchal et al., 2016; Toum et al., 2016). COR appears to contribute more significantly to *Pst* virulence at night in *Arabidopsis thaliana* (hereafter, *Arabidopsis*) (Panchal et al., 2016), suggesting either a time of day and/or an effect of light on bacterial interaction with stomata.

In contrast to early events in infection that promote stomatal opening, pathogens appear to induce stomatal closure in later stages (Hu et al., 2022; Roussin-Léveillée et al., 2022). A frequent feature of pathogen infection in plants is the creation of an aqueous environment in the plant apoplast, visually noticeable as water-soaked lesions, which is crucial for virulence (Xin et al., 2016). *Pst* encodes two highly conserved effector proteins, HopM1 and AvrE1, that induce water-soaking lesions (Hu et al., 2022; Roussin-Léveillée et al., 2022; Xin et al., 2016). Notably, these water-soaking effectors stimulate host abscisic acid (ABA) biosynthesis and signaling pathways to induce stomatal closure, resulting in a loss of transpiration and accumulation of water in the apoplast (Hu et al., 2022a; Roussin-Léveillée et al., 2022).

Stomatal status is strongly influenced by diurnal light cycles, and light also affects plant responses to pathogens (Carvalho & Castillo, 2018; Shah et al., 2021). Moreover, SA regulates phytochromes (PHY) proteins, which act as photoreceptors (Shah et al., 2021), and PHYA and PHYB have been reported to be required for the induction of SA-responsive genes (Genoud et al., 2002; Karpinski et al., 2003). PHYTOCHROME-INTERACTING FACTORs (PIFs) are rapidly degraded during red-light perception by PHYs and are also involved in light-dependant modulation of immune signaling (Gangappa & Kumar, 2018). Furthermore, functional circadian cycles are required to respond to many abiotic and biotic stresses (Roeber et al., 2022), including responses to infection (Karapetyan & Dong, 2018). Indeed, plant susceptibility to pathogens varies, depending on the time of day of infection (Griebel & Zeier, 2008; Zhang et al., 2013). It has been reported that plants are less susceptible in the subjective morning because defense responses are regulated by the circadian clock (Bhardwaj et al., 2011). Accumulation of SA has been reported to be circadian-gated, with levels increasing throughout the day and decreasing throughout the night (Zheng et al., 2015). Moreover, the circadian regulators CCA1 and LHY play key roles in defense against *Pst*, as they promote stomatal closure in early defense responses in a SA-dependant manner (Zhang et al., 2013). However, as circadian cycles are tightly linked to light perception, it is important to determine if a given phenomenon is related to the presence of light or to circadian mechanisms *per se*.

In this study, we report a novel role for light in modulating plant defense responses. Constant light treatment prevents bacterial infection by abrogating the ability of *Pst* to induce stomatal closure, thereby inhibiting the accumulation of water in the apoplast. This effect is due to a potentiation of SA signaling, which induces a stomatal opening activity that dominates over ABA. In addition to highlighting the interplay between abiotic and biotic stress, our results shed light on a physiological mechanism by which SA contributes to plant defense against pathogens.

## Results

### Constant light confers protection against *Pseudomonas syringae*

To study the influence of light on the outcome of a bacterial infection in plants, we evaluated the virulence of *Pst* on *Arabidopsis thaliana* Col-0 wild-type (WT) plants under different light regimes. Plants were grown under 12 hours light/dark cycles before being challenged with *Pst* and placed under either constant light (LL), 12 hours light/dark (LD) or constant dark (DD) regimes for three days. We found that *Pst-*infected *Arabidopsis* plants kept in LL showed a drastic reduction in disease symptoms compared to those kept in LD or DD over the course of the infection (Fig. 1a). This observation was reflected in bacterial titers, as plants kept under LL accumulated 10-fold less *Pst* compared to other conditions (Fig. 1b). Interestingly, constant light had no additional protective effect against *Pst* DC3000 *hrcC*^*-*^, which lacks a T3SS, suggesting that this regime affects effector protein actions (Fig. 1b, c).

**Figure 1:**
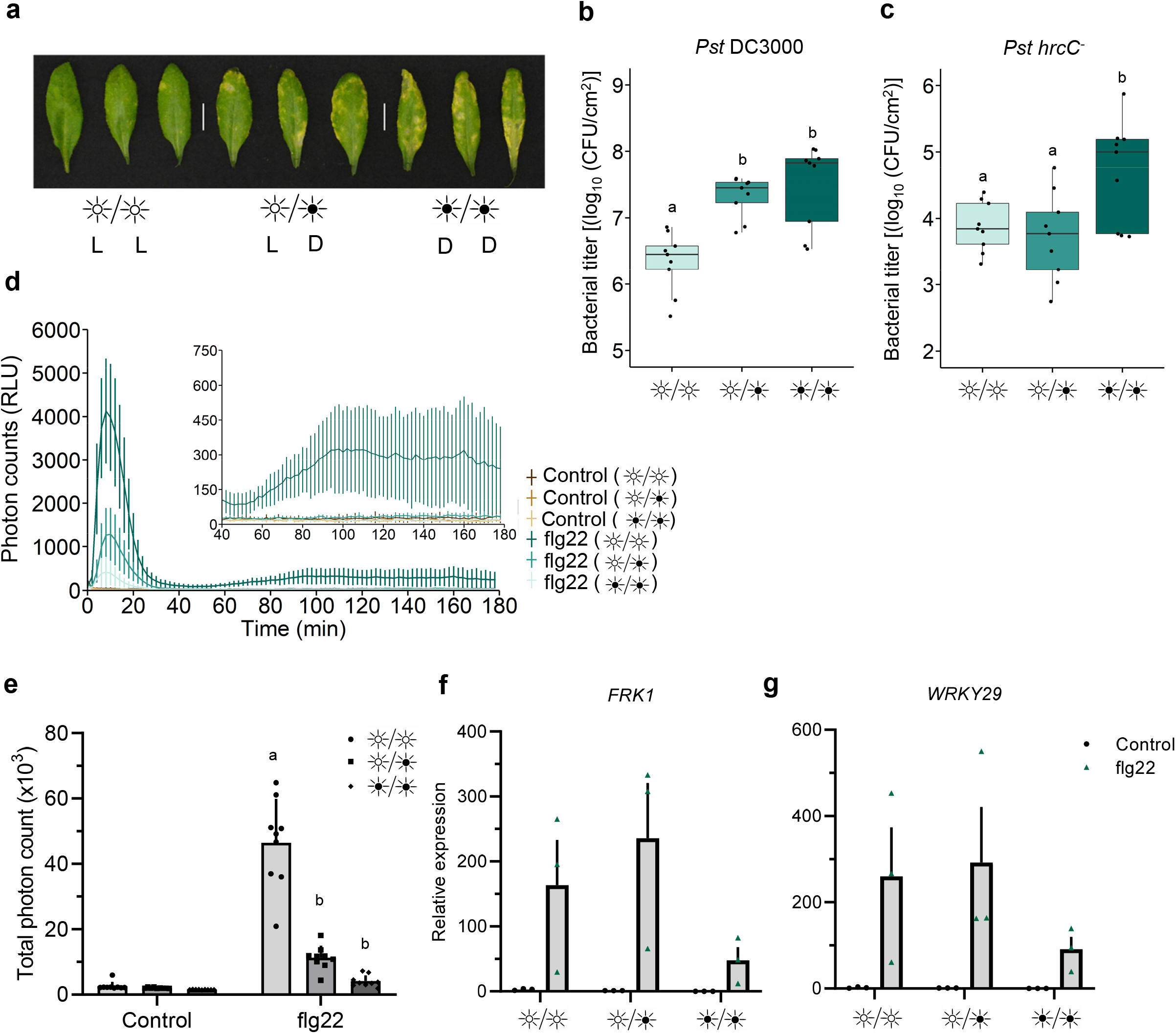
Constant light confers protection against *Pst*. **a**, Disease phenotypes in *Arabidopsis* leaves syringe-infiltrated with *Pst* DC3000 (1×10^5^ CFU/ml). WT Plants were either kept under constant light (LL), 12 hours light/dark (LD) or constant dark (DD) light cycles for three days after inoculation. **b-c**, Bacterial titers from WT leaves syringe-infiltrated with *Pst* DC3000 (**b**) or *Pst hrcC*^*-*^ (**c**) at three days n post-infection (dpi) under light regimes described in **a. d**, Apoplastic oxidative burst triggered by 1 μM flg22 in WT *Arabidopsis* leaf punches. Leaf punches were kept in water under LL, LD or DD for 24 hours prior to stimulation with flg22. Values are averages ± se (n=27). **e**, Total photon counts measured over 180 minutes from the experiments in **d. f-g**, Expression levels of the early PTI marker genes *FRK1* (**f**) and *WRKY29* (**g**) treated with 1 μM flg22 in plants that were previously placed in the indicated light setting and measured by qRT-PCR. Samples were kept under these light setting for 6 hours after control or flg22 treatment after which they were harvested. Different letters indicates statistically significant differences, p < 0.05, ANOVA (**b, c**) or p < 0.0001, ANOVA followed by Tukey’s range test.

Circadian rhythms and light regulation have been shown to impact bacterial virulence (Griebel & Zeier, 2008; Wang et al., 2011; Zhang et al., 2013). Therefore, we evaluated whether plants subjected to different light regimes for a 24-hour period displayed irregular modulation of immunity following treatment with the immunogenic peptide flg22. We first explored the production of apoplastic reactive oxygen species (aROS), a commonly used early marker of immune responses. We found that plants subjected to LL for 24 hours prior to treatment with flg22 produced significantly more aROS than plants kept under LD or DD regimes (Fig. 1d, e). Notably, under LL, *Arabidopsis* plants displayed a moderate, but persistent second wave of aROS, as opposed to other light conditions (Fig. 1d). In contrast, plants that were subjected to DD treatment for a single day produced much less aROS in response to flg22, compared to plants kept under LD conditions (Fig. 1e). Hence, light appears to prime *Arabidopsis* plants to produce enhanced and more sustained aROS after immune elicitation, potentially contributing to disease resistance.

We evaluated the expression of two early PTI-responsive genes, *FRK1* and *WRKY29*, to assess early PTI responsiveness. Plants were subjected to the three different light regimes for 24 hours before being challenged with flg22 for four hours. Although not significantly altered, these genes were less expressed in DD compared to LD or LL (Fig. 1f, g). Interestingly, constant light by itself was able to modulate both FLS2 and RBOHD expression levels in the absence of flg22 (Fig. S1a, b). Contrary to this observation, *FLS2* and *RBOHD* expression levels were not upregulated upon flg22 treatment under constant dark (Fig. S1a, b). These results suggests that light perception and/or signaling is critical for the establishment of a proper early PTI response.

### *Pseudomonas syringae* requires darkness to induce water-soaking lesions in its host

The virulence of *Pst* is greatly enhanced by its ability to induce water-soaking lesions by closing stomata under high humidity conditions (Hu et al., 2022; Roussin-Léveillée et al., 2022). As light modulates stomatal aperture (Matthews et al., 2020), we questioned whether a pathogen’s ability to induce stomatal closure could be affected by diurnal light oscillations. We observed that, under LL conditions, *Pst* could no longer induce water-soaking lesions in *Arabidopsis* leaves, in contrast to LD or DD (Fig. 2a). The latter was not due to differences in levels of bacteria, as inoculations were carried with a high concentration of bacteria (1 × 10^8^ CFU/ml) (Fig. 2b). Consistent with the lack of water soaking, we found that stomata of inoculated *Arabidopsis* plants kept under LL conditions were still fully open 24 hours post inoculation (Fig. 2c). In contrast, stomata of inoculated plants kept under LD conditions were closed, as expected, as were those of plants kept under DD conditions (Fig. 2c).

**Figure 2:**
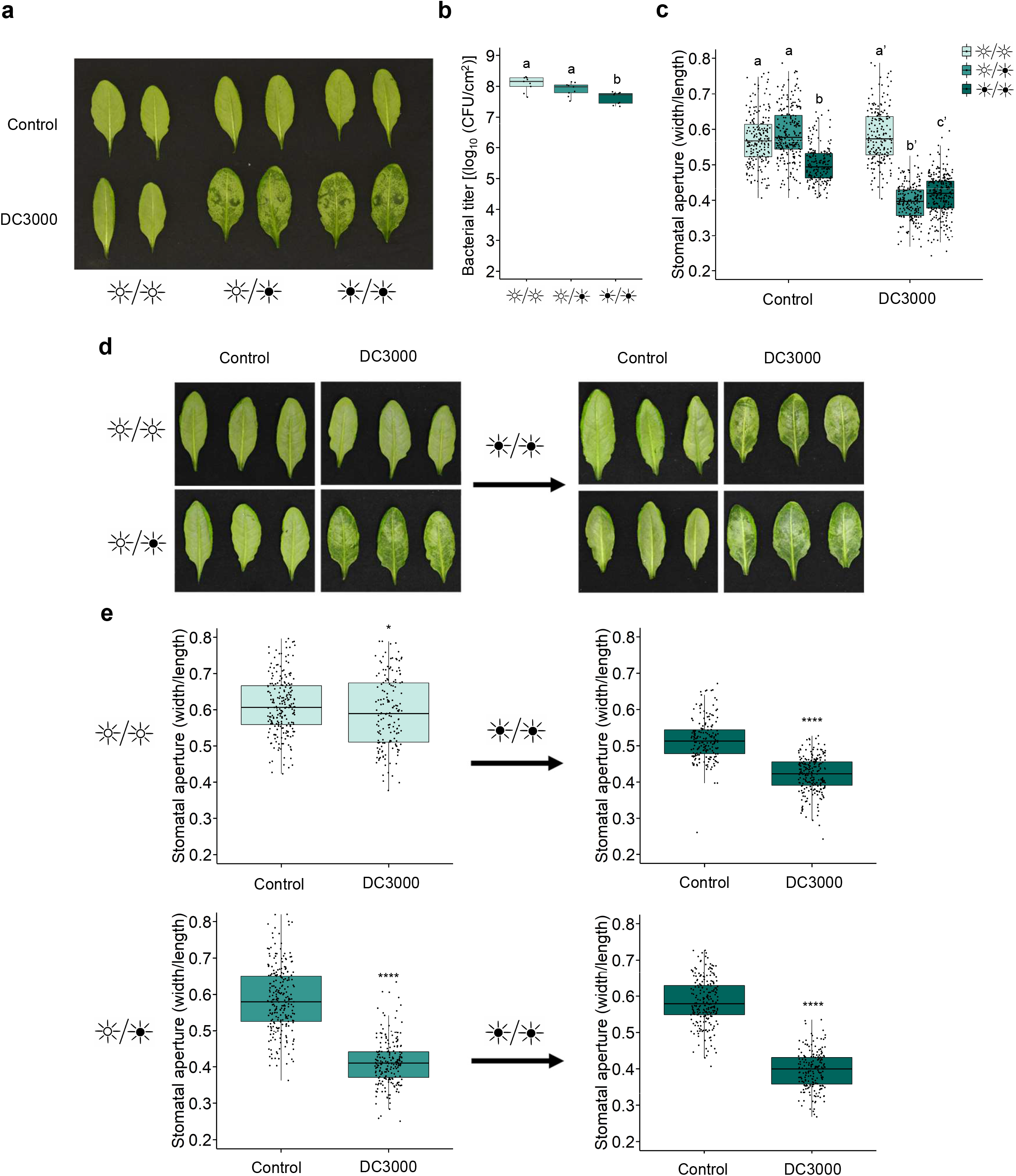
*Pst* requires darkness to induce stomatal closure and water-soaking. **a**, Water-soaking phenotypes in *Arabidopsis* leaves syringe-infiltrated with *Pst* DC3000 (1 × 10^8^ CFU/ml) at 24 hpi under LL, LD or DD conditions. **b**, Bacterial titers from plants described in **a. c**, Stomatal aperture measurements in plants described in **a** (n > 150 stomata) by epifluorescent microscopy. **d**, Water-soaking phenotypes in *Arabidopsis* leaves that were syringe-infiltrated with *Pst* DC3000 (1 × 10^8^ CFU/ml) at 24 hpi under LL or LD conditions (left panel) and in plants treated in the same way followed by transferred to DD for 6 hours (right panel). **e**, Stomatal aperture measurements from leaves treated as in **d** (n > 150 stomata). Different letters indicate statistically significant differences, p < 0.05, ANOVA (**b**) or Kruskal-Wallis test (**c**). Asterisks indicate statistically significant differences compared to control, * p < 0.05, **** p < 2.2 × 10^−16^, Student’s t test (**e**, upper right panel) or Wilcoxon-Mann Witney test (**e**, upper left, lower right and lower left panels).

To consolidate our findings, we designed an experiment where *Arabidopsis* plants were infected with *Pst* and either kept under LL or LD conditions for the first 16 hours of the infection. At this timepoint, LD, but not LL treated plants underwent water soaking (Fig. 2d). Plants were then placed in darkness to determine if water soaking could be reappearing. Indeed, we found that the water-soaking phenotype could be fully restored after keeping LL treated plants in darkness for six hours (Fig. 2d). Stomatal behavior analysis further supported the observation that constant light prevents *Pst* from inducing stomatal closure, but that this phenomenon is swiftly reversed once under dark conditions (Fig. 2e).

*Pst* manipulates its host ABA machinery via secretion of its conserved effectors HopM1 and AvrE1 to induce stomatal closure leading to water-soaking (Hu et al., 2022; Roussin-Léveillée et al., 2022). We designed experiments to test whether a deficiency in apoplast hydration was the main reason behind reduced *Pst* aggressivity under LL. Since the *Arabidopsis aba2-1* mutant was shown to prevent *Pst*-induced water-soaked lesions (Hu et al., 2022; Roussin-Léveillée et al., 2022), we compared the bacterial levels of these plants infected under LL and LD conditions. We found that, while bacterial population levels were affected in WT plants between LL and LD conditions, they were unaltered in *aba2-1* plants (Fig. S2). Moreover, under LL, *Pst* grew to levels similar to *Pst hopM1-/avrE1-* (*h-/a-*), which cannot induce water-soaking (Fig. S3). Furthermore, the pathogen’s ability to manipulate ABA appeared unaffected as transcript levels of the ABA biosynthesis and signaling marker genes, *NCED3* and *RD29A*, respectively, in WT infected plants were similar under all light conditions (Fig. S4). These results suggest that the lack of apoplastic fluid could at least partly explain the growth deficiency of *Pst* under LL. Thus, it appears that pathogens might benefit from natural light cycles to establish a favorable microhabitat inside their hosts.

### Salicylic acid is required to prevent effector-mediated induction of water-soaking under continuous light

Salicylic acid is a key immune component and antagonizes the molecular multiple actions driven by abscisic acid signaling (de Torres Zabala et al., 2009; Meguro & Sato, 2014; Moeder et al., 2010). However, how SA and ABA signaling interact with each other with respect to stomatal control during late phases of pathogen infection is unknown. We investigated whether plants that were subjected to a constant light cycle for 24 hours would be altered in salicylic acid (SA) signaling or accumulation. First, we analysed expression levels of SA responsive-genes, *ICS1* (*ISOCHORISMATE SYNTHASE 1*) and *PR1* (*PATHOGENESIS-RELATED PROTEIN 1*), which are often used to assess SA biosynthesis and responsiveness, respectively. We observed that the expression levels of these two genes were increased under LL conditions relative to LD or DD regimes, although the increase of *ICS1* relative expression was not significant (Fig. 3a, b). Consistent with this, we observed, using ultra performance liquid chromatography-mass spectrometry (UPLC-MS), that upon treatment with flg22, plants kept under LL conditions showed an increase in SA levels compared to LD and DD treated plants (Fig. 3c). As such, constant light appears not only to modulate early PTI responses, but also SA-related responses.

**Figure 3:**
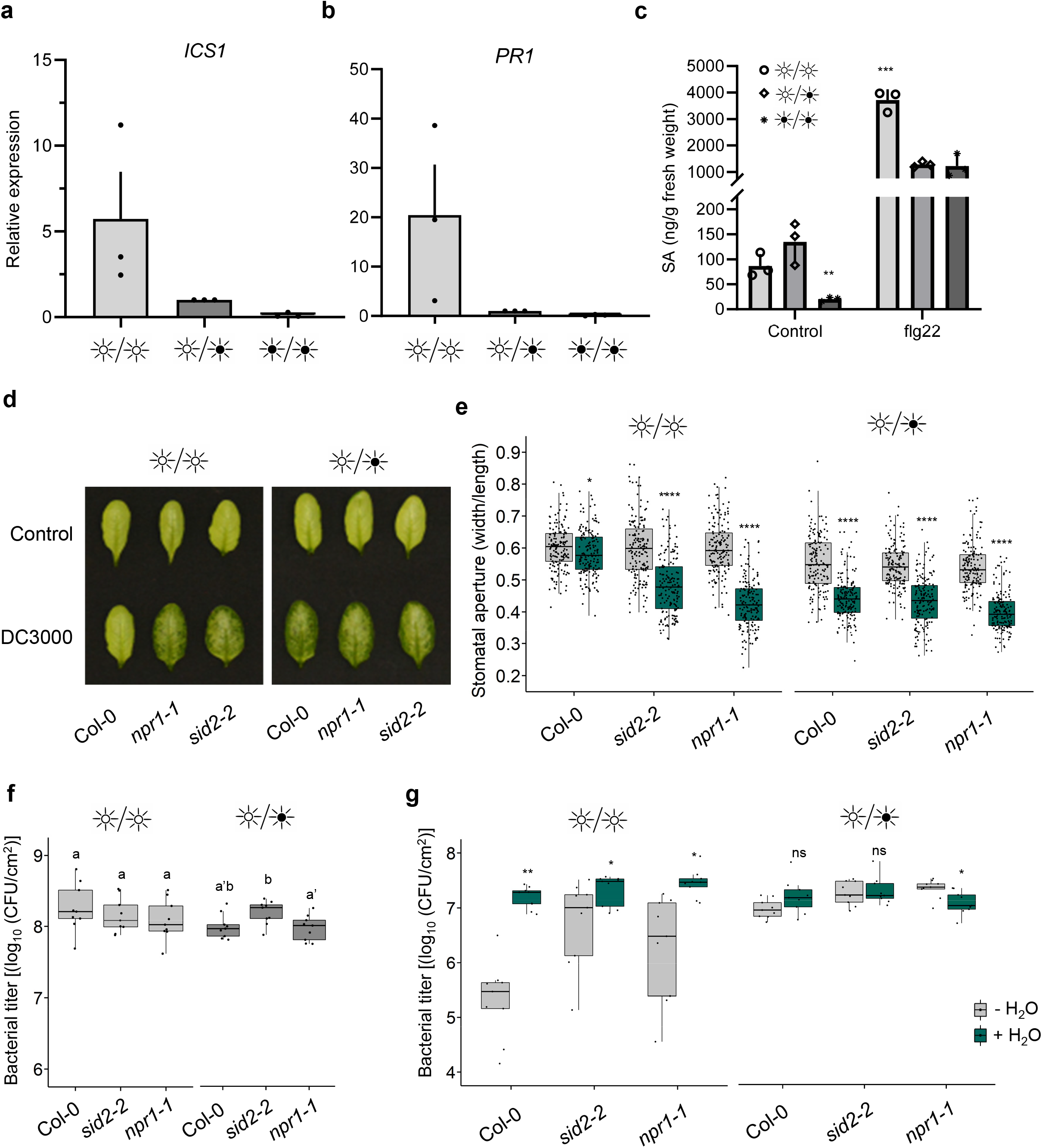
Salicylic acid is required for preventing stomatal closure and disease progression under constant light. **a-b**, Expression levels, measured by qRT-PCR, of SA biosynthesis (*ICS1*) and signaling (*PR1*) marker genes in WT *Arabidopsis* plants treated with 1 μM flg22 after having previously been placed in the identified light settings. Samples were harvested at 24 hpi. **c**, Quantification of SA in WT *Arabidopsis* plants mock inoculated (control) or inoculated with 1 μM flg22 under the same experimental settings as in **a** measured by UPLC-MS at 24 hpi. **d**, Water-soaking phenotype in *Arabidopsis* WT (Col-0), *sid2* and *npr1* mutant plants syringe-infiltrated with *Pst* DC3000 (1 × 10^8^ CFU/ml) at 24 hpi under LL, LD or DD. **e**, Stomatal aperture measurements from infected leaves displayed in **d** (n > 150 stomata). **f**, Bacterial titers from *Arabidopsis* WT (Col-0), *sid2* and *npr1* mutant plants infected with *Pst* DC3000 (1 × 10^8^ CFU/ml) at 24 hpi. **g**, Bacterial titer of *Arabidopsis* WT (Col-0), *sid2* and *npr1* mutant plants infected with *Pst* DC3000 (1 × 10^5^ CFU/ml) in which infiltrated leaves were allowed to return to a pre-infiltrated state (-H_2_O) or in plants that were immediately domed following infiltration (+H_2_O). Bacterial titers were measured at 3 dpi. Asterisks indicate statistically significant differences compared to control, ns = non-significant, * p < 0.05, ** p < 0.001, **** p < 2.2 × 10^−16^, ANOVA followed by Tukey’s range test compared to LD (**c**), Student’s t test (**e, f**) or Wilcoxon-Mann Witney test (**e, g**).

In agreement with our findings, a recent report revealed that SA-related responses are primed by photoperiodic stress (Cortleven et al., 2022). Considering that SA can antagonize ABA responses, we investigated whether SA is necessary for light-mediated prevention of water-soaked lesions. We inoculated *Arabidopsis* mutants, *npr1-1* and *sid2-2*, which are compromised in SA signaling and biosynthesis, respectively, with *Pst* under different light conditions. Interestingly, we observed water-soaking in the inoculated leaves of *npr1-1* and *sid2-2* mutant plants at 24 hours post-infection (hpi) under both LD and LL conditions, whereas wild-type plants showed water-soaking only under LD conditions (Fig. 3d). Consistent with these results, we observed that *Pst* inoculation induced stomatal closure at 24 hpi in all genotypes under LD conditions (Fig. 3e). Furthermore, apoplastic hydration was elevated in LL in SA-related mutants, but not in Col-0 under LL (Fig. S5). However, constant light did not hamper the ability of the pathogen to induce stomatal closure in the *npr1-1* mutant, or (to a lesser degree) in the *sid2-2* mutant, as it does in WT plants (Fig. 3e). These altered responses were not due to differences in bacterial densities, as bacterial titers did not differ in mutant versus wild-type plants at this time point (Fig. 3f).

The impact of SA on plant responses to infection may be due to multiple effects. Given the effect of SA on water-soaking and stomatal control (Fig. 3d, e), we tested if the effect of SA might be related to the establishment of an aqueous environment, which is beneficial to bacteria. To do so, we employed an inoculation method wherein plants are immediately placed in a >95% humidity environment after inoculation without allowing liquid from the inoculum to evaporate. Under these conditions, leaves remain saturated with water, mimicking at least in part the phenomenon of water-soaking. Artificial induction of water soaking had no effect on bacterial growth of either wild-type or mutant plants under LD conditions (Fig. 3g). Keeping plants under LL conditions resulted in a dramatic decrease in bacterial growth in WT plants, while this growth defect was partially rescued in *sid2-2* and *npr1-1* mutants (Fig. 3g). Importantly, artificial water-soaking resulted in a dramatic increase of bacterial growth under LL conditions, with both WT and mutant plants showing bacterial levels similar to LD grown plants (Fig. 3g). These results indicate that the induction of resistance induced by constant light requires SA signalling and that the effects of SA are, at least in part, related to its effects on inhibiting stomatal closure. This, in turn, inhibits the pathogen’s ability to create an ideal aqueous environment in the apoplast.

### BTH suppresses *Pst*-induced water-soaking lesions

Benzothiadiazole (BTH) is an analog of SA and, like SA, triggers systemic acquired resistance in plants and confers protection against *Pst* (Görlach et al., 1996; Kim et al., 2022). Since SA biosynthesis and signaling are essential for preventing water-soaking under LL (Fig. 3d), we explored the potential of using BTH to prevent this phenomenon. Plants that were pre-treated with BTH 24 hours prior to a *Pst* challenge did not display water-soaking lesions, in contrast to control-treated plants (Fig. 4a). Most importantly, BTH prevented *Pst*-induced water-soaking lesions under a LD regime, in which water-soaking lesions are normally visible. Consistent with this result, *Pst* did not induce stomatal closure in LD-grown plants pre-treated with BTH (Fig. 4b). These effects were independent of the number of bacteria, as bacterial titers in leaves did not differ between treatments at the time of stomatal evaluation (Fig. 4c).

**Figure 4.**
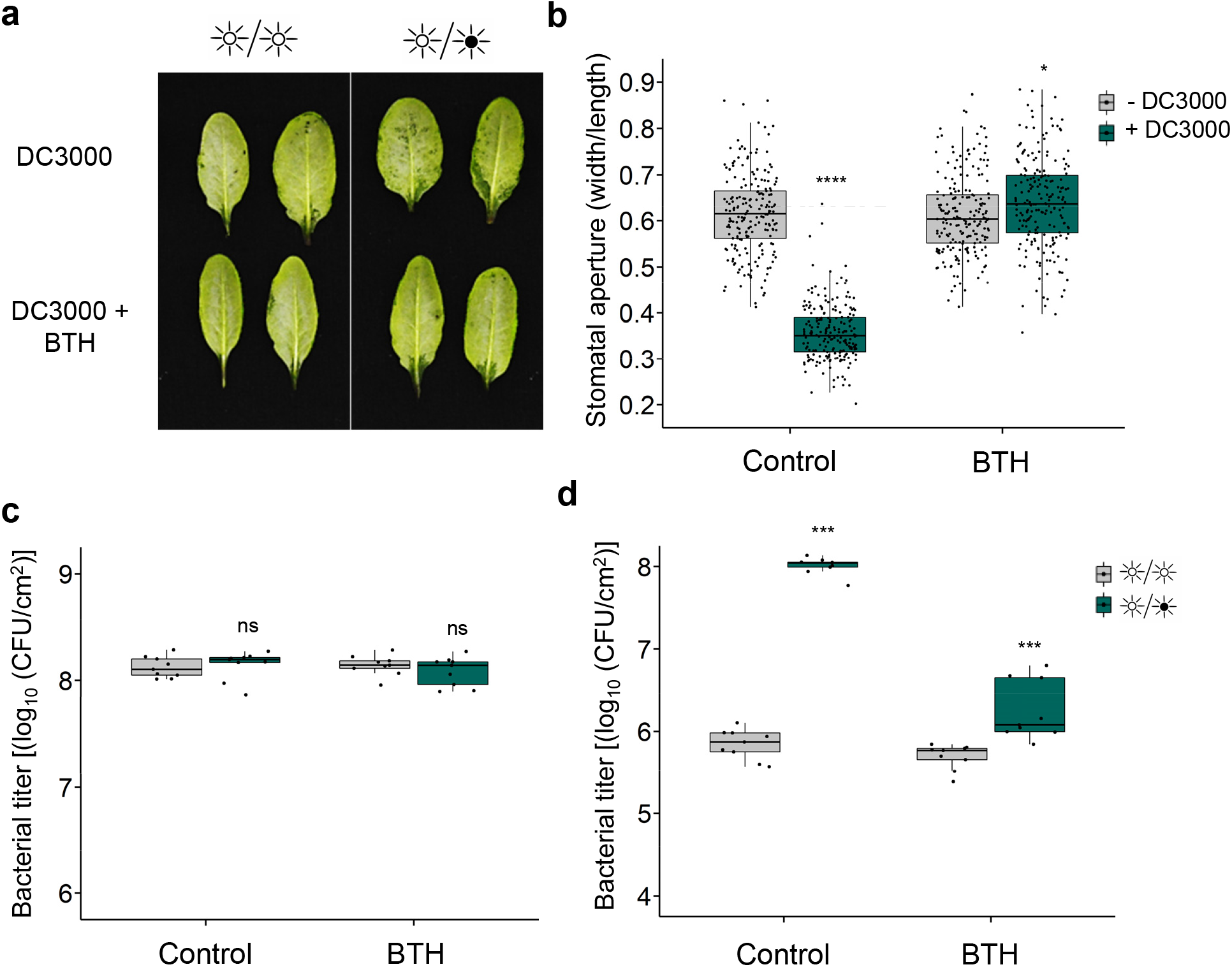
BTH provides protection against bacterial-induction of water-soaking. **a**, Water -soaking phenotypes in WT *Arabidopsis* infiltrated with DMSO (0.1%) or BTH (50 μM) prior to syringe-infiltration with *Pst* DC3000 (1 × 10^8^ CFU/ml) 24 hours later. Photos were taken 24 hours post infection. **b**, Stomatal aperture in WT *Arabidopsis* treated as in **a** and in which MgCl_2_ (10 mM; - DC3000) or *Pst* (1 × 10^8^ CFU/ml; + DC3000) was syringe-filtrated. Plants were left under light/dark conditions and stomatal apertures assessed at 24 hours post infiltration (hpi). **C**, Bacterial titers from plants treated in **a** measured 24 hpi. **d**, Bacterial titers from *Arabidopsis* leaves that were infiltrated with DMSO (0.1%) or BTH (50 μM) 24 hours prior to being challenged with *Pst* DC3000 (1 × 10^5^ CFU/ml). Plants were placed under the indicated light conditions after infection. Bacterial titers were assessed at 3 days post infection and plants were kept under the indicated light conditions throughout the infection process. Asterisks indicate statistically significant differences compared to control, ns = non-significant, *** p < 0.0001, Student’s t test (**c**) or Wilcoxon-Mann Witney test (**d**).

We next tested the ability of BTH to mitigate a *Pst* infection under either LL or LD in locally pre-treated leaves. Interestingly, levels of bacteria in control or BTH pre-treated leaves were similar under constant light, suggesting that BTH does not induce more resistance than a photoperiodic stress alone (Fig. 4d). However, a clear difference was observed between control and BTH pre-treated plants under LD conditions (approx. 100-fold less bacteria). At the same time, LD-grown plants pretreated with BTH showed a protection close, albeit statistically significantly different to that accorded by growing under LL, (5.69±0.15 CFU/cm^2^ (LL); 6.25±0.36 CFU/cm^2^ (LD)) (Fig. 4d). These results suggest that BTH provides protection similar to that provided by a constant light treatment, and further indicates that the effects of constant light in this context are due to the actions of SA.

### Constant light treatment allows plants to recover from bacterial infection

Given the role of light in preventing disease symptoms and pathogen growth, we evaluated whether constant light could mitigate the progression of the pathogen in plants that were already infected by a pathogen. To test this, plants were inoculated and left under a LD cycle for a day to allow the pathogen to establish an infection. Plants were then transferred to LL conditions for two days, then again to a LD cycle for two additional days. Control plants were similarly inoculated but kept under a LD cycle for the whole duration of the experiment (Fig. 5a). No disease symptoms were observed on plant leaves that had been subjected to the two-day LL treatment (Fig. 5b). Bacterial titer assays were also performed and a decrease in bacterial load was observed when plants were kept under LL conditions compared to plants kept under LD conditions for the whole experiment (Fig. 5c). These results suggest that constant light treatment allows plants to recover from a *Pst* infection.

**Figure 5:**
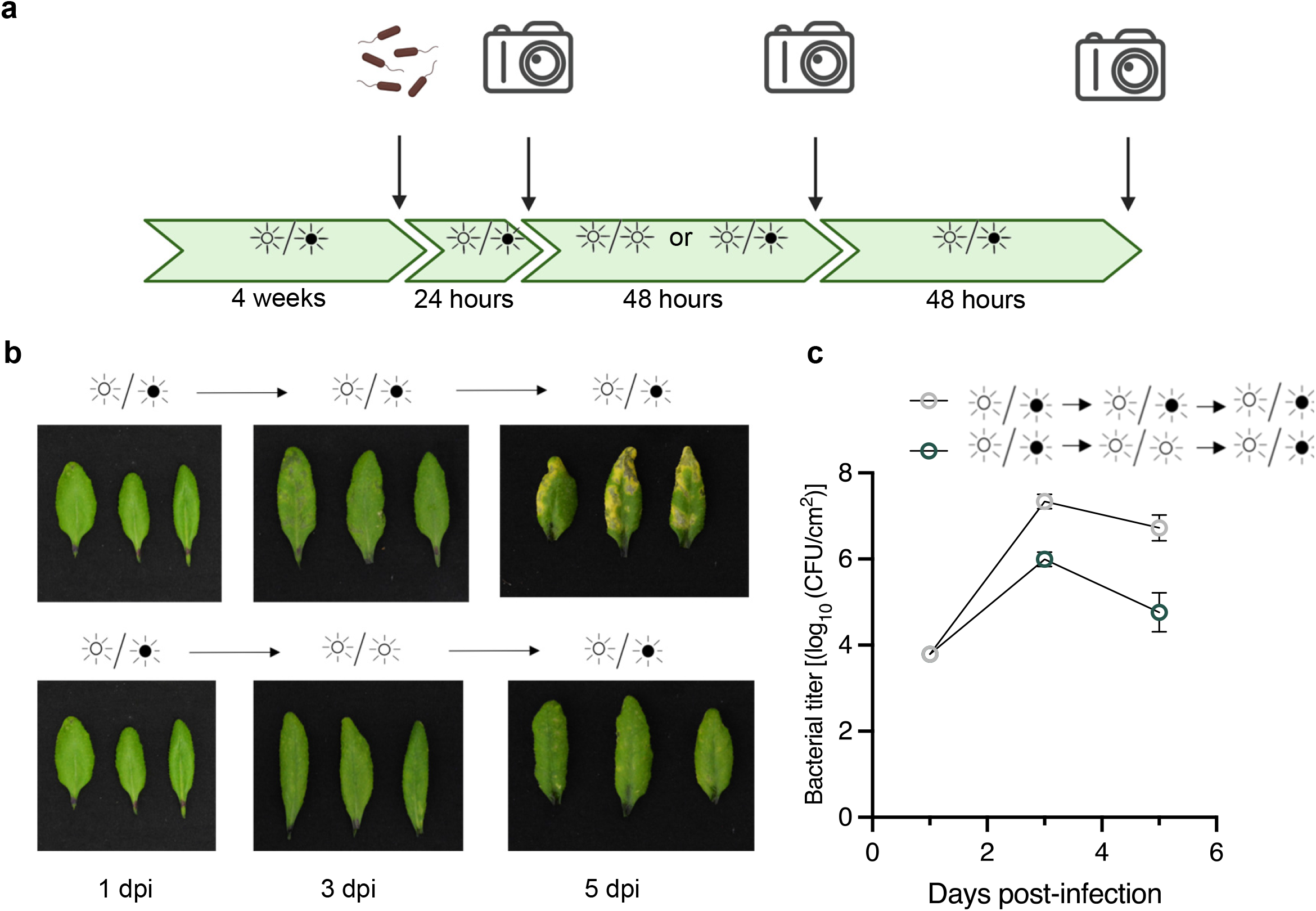
Using constant light to prevent disease development. **a**, Schematic diagram representing treatments and phenotype assessment of light-mediated disease prevention. **b**, Disease symptoms in *Arabidopsis* plants infected with 1 × 10^5^ CFU/ml under different light settings, as, indicated. **c**, Bacterial titers from plants shown in **b** and harvested at the indicated timepoints.

### PIFs modulate disease resistance against *Pst* under constant light potentially through SA signaling

We investigated the implication of light photoreception and signaling in susceptibility to bacterial infection under LL or LD regimes. For this, we used the *Arabidopsis* blue-light insensitive (*blus1*) mutant as well as a quadruple mutant for phytochrome-interacting factors (*pifq*), which is responsible for integrating red and far/red light signaling. The *Arabidopsis blus1* mutant behaved similarly to wild-type plants in terms of water-soaking under LD and LL (Fig. 6a). However, *pifq* mutants displayed a reestablishment of water-soaked lesions 24 hours after inoculation with *Pst* (Fig. 6a). These observations were consistent with measurements of stomatal aperture, wherein only *pifq* mutants infected with *Pst* had closed stomata under constant light (Fig. 6b). Interestingly, all three genotypes showed similar increases of induction of expression of *ICS1* upon infection with *Pst*, with somewhat higher levels in LL versus LD (Fig. 6c). However, whereas WT and *blus1* plants showed much higher induction of *PR1* expression in LL versus LD, *pifq* mutant plants kept under LL conditions showed *PR1* induction similar to WT plants grown in a LD regime (Fig. 6d). Thus, the levels of *PR1* induction and stomatal closure in LL-grown *pifq* plants resemble those of LD-grown WT plants, further underlining the link between SA signaling, stomatal closure, and water-soaking. These results also suggest that PIFs might be involved in strengthening the SA module during constant light after immune elicitation.

**Figure 6:**
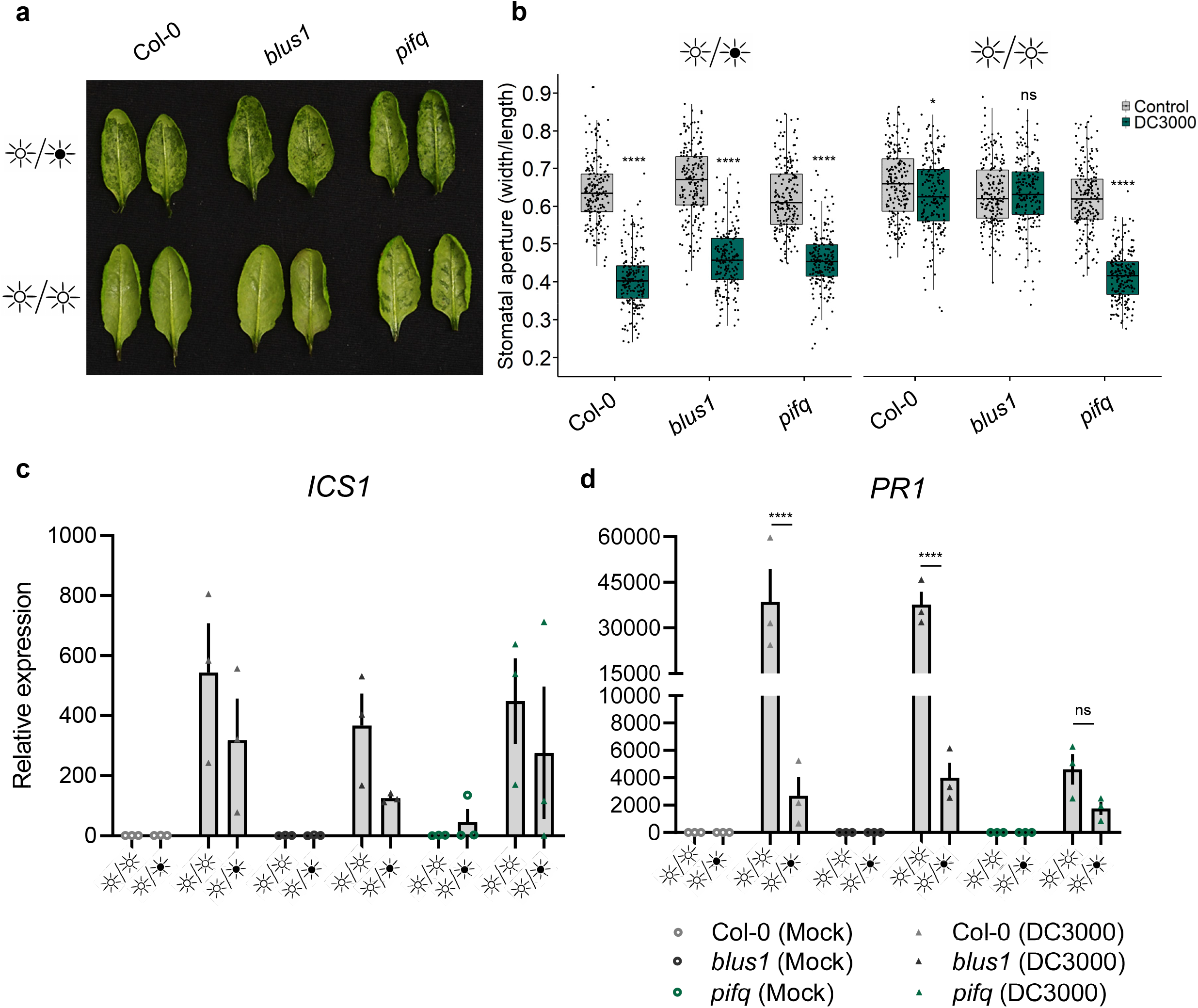
Potential implication of PIFs in preventing bacterial induction of water-soaked lesions under ight. **a**, Water-soaking phenotypes of *Arabidopsis* WT (Col-0), *blus1* and *pifq* mutant plants syringe-infiltrated with *Pst* 3000 (1 x10^8^ CFU/ml) under the indicated light settings. Photos were taken at 24 hours post infiltration (hpi). **b**, Stomatal aperture measurements from infected leaves displayed in **a** (n > 150 stomata). **c-d**, Expression levels, assessed by qRT-PCR, of *ICS1* (c) and *PR1* (d) marker genes in *Arabidopsis* WT (Col-0), *blus1* and *pifq* mutant plants infected with *Pst* DC3000 (1 × 10^8^ CFU/ml) or with control solution (MgCl_2_ 10 mM) in plants that were placed in the identified light settings. Samples were harvested at 24 hpi. Asterisks indicate statistically significant differences compared to control, ns = non-significant, * p < 0.05, **** p < 2.2 × 10^−16^, Wilcoxon-Mann Witney test (**b**) or **** p < 0.0001, Student’s T-test (**d**).

## Discussion

Induction of a water-soaked apoplast is crucial for disease development in plants (Aung et al., 2018; Peng et al., 2019; Xin et al., 2016; D. Zhang et al., 2019). In this study, we provide evidence that diurnal light cycles are intimately linked with the ability of a bacterial pathogen to establish an aqueous apoplast. We show that a period of darkness is required to induce water-soaking and that constant light prevent induction of stomatal closure mediated by water-soaking effectors. Light counteracts the effects of ABA and prevents stomatal closure through potentiation of SA responses. Furthermore, BTH, a SA analog, was sufficient to prevent bacterial-induction of water-soaking lesions. Considering how common the induction of water-soaked lesions is as an initial step in microbial pathogenesis (Aung et al., 2018), finding non-invasive and easily applicable solutions to prevent them is of great practical interest.

In studies of the circadian clock machinery plants are often placed in constant light to study oscillations in gene regulation and other phenomena (Velez-Ramirez et al., 2011). As such, much of our understanding of circadian-gated regulation of phytohormone networks rely heavily on data obtained under constant light regimes (Li et al., 2018; Zheng et al., 2015; Zhou et al., 2015). However, light can affect molecular processes independently of circadian mechanisms. Indeed, *Arabidopsis* plants with mutations in key circadian cycle-regulating genes are protected against infection when grown under LL conditions compare to LD. Importantly, the degree of protection is similar to that seen with WT plants (Fig. S6a, b), suggesting that the effects of constant light are not dependant on the circadian cycle.

Our data suggest that, under constant light regimes, plant immune responses are augmented or primed. Indeed, apoplastic ROS bursts following flg22 perception are more intense under LL conditions and less intense under DD (Fig. 1d). These results are consistent with the elevated levels of expression of *FLS2, RBOHD, ICS1* and *PR1* under LL conditions prior to flg22 treatment (Fig. S1, 3a, b). However, the expression of early PTI-responsive genes was not significantly altered between light conditions prior to, or after, treatment with flg22 (Fig. 1f, g). Nonetheless, consistent with increased aROS production, plants grown under LL conditions appear to respond with greater intensity to flg22 in terms of SA accumulation (Fig. 3c). How light affects the responsiveness to immune elicitors remains to be investigated. However, our results showing that it affects aROS production and SA-related responses, but not early PTI responses, suggest that constant light affects specific modules in plant immunity. At the same time, our results are consistent with a recent study showing that, under photoperiodic stress, the *Arabidopsis* transcriptional signature resembles that of a response to pathogens (Cortleven et al., 2022). Notably, SA signatures were predominant in plants kept under constant light and were reduced in dark-grown plants.

It has been reported that pathogen aggressiveness is increased in infections occurring in the shade (Roberts & Paul, 2006). However, whether this involves host stomatal responses and is related to pathogen-induced water-soaking lesions has not been investigated. Diurnal light changes affect stomatal aperture, where stomata are open during the day and partially close during the night (Yang et al., 2020). As such, a synergistic interaction may occur in which the mechanism(s) behind dark-induced stomatal closure, while not well defined, may act as facilitators for effector-mediated induction of stomatal closure. It is important to note that, in infected plants, light-induced inhibition of stomatal closure and water-soaking can be swiftly reversed. That is, non-water-soaked infected plants with LL and SA-dependent open stomata undergo water-soaking after a four-hour dark treatment (Fig. 2d, e). This suggests that dark conditions may be a prerequisite for the induction of water-soaking. Current understandings on the mechanism behind darkness-induced stomatal closure suggest that ABA signaling and metabolism are important for the response speed, but not essential for closure, suggesting ABA-dependent and independent mechanisms (Pridgeon & Hetherington, 2021). Thus, we speculate that the balance in the antagonism between ABA and SA and stomatal aperture may be affected by additional mechanisms that depend on light (Fig. 7). This could include regulation of ABA levels, which increase substantially within a four-hour period of darkness (Weatherwax et al., 1996).

**Figure 7:**
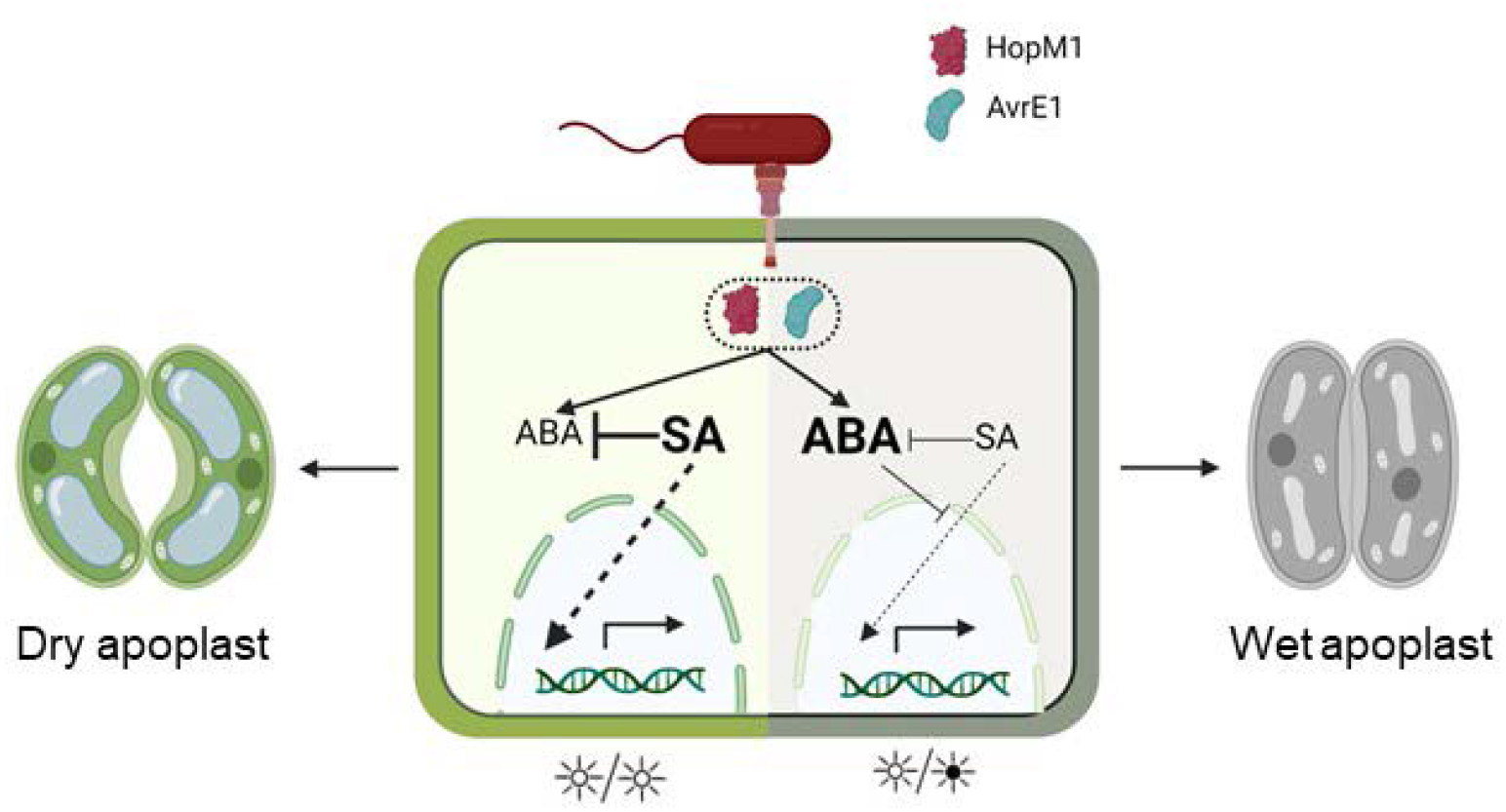
Model for light-induced disease resistance in plants.

Water-soaking effectors of *Pst* modulate ABA-related responses (Hu et al., 2022; Roussin-Léveillée et al., 2022). The induction of expression of ABA biosynthesis and signaling pathways by *Pst* was unaltered between all light regimes tested (Fig. S4). This suggests that the reported SA antagonism of ABA-related responses may be downstream of ABA responsive targets (de Torres Zabala et al., 2009; Meguro & Sato, 2015; Moeder et al., 2010). Our observations are reminiscent of a previous report showing that SA accumulation antagonizes ABA responses, including stomatal closure, even if ABA over accumulates and certain ABA-responsive genes are activated at the transcriptional level (Moeder et al., 2010). Whether SA-mediated antagonism of ABA responses is solely responsible for the pathogen’s inability to induce an aqueous apoplast remains to be investigated.

ABA is a major susceptibility factor to most known ABA producing or ABA inducing plant pathogens (Lievens et al., 2017). In addition to *Pst*, exposure to photoperiodic stress leads to enhanced disease resistance to the hemi-biotrophic fungus *M. oryzae* and the necrotrophic fungus *B. cinerea* present enhanced disease resistance (Cagnola et al., 2018; Shimizu et al., 2021). Interestingly, these two fungal pathogens produce ABA and induce water-soaking lesions early on during the infection process. Whether water-soaking contributes to fungal pathogenicity is not clear, but ABA was found to be a key virulence component for *M. oryzae* and *B. cinerea* (Audenaert et al., 2002; Spence et al., 2015). It is thus interesting to speculate that the reduced virulence of these fungal pathogens in plants exposed to photoperiodic stress is caused by their inability to create an aqueous apoplast as well. Therefore, exposing plants to constant light for a short amount of time could be a very impactful strategy to reduce infections from pathogens from different kingdoms of life.

We are still in the dark about the specific wavelengths of light that affect the defense responses described herein. However, it has been reported that red light treatment improves resistance against *Pst* and that this treatment increased SA signaling as well as transcription of genes involved in redox homeostasis, such as *RBOH* and *GSTs* (Yang et al., 2015). Our data provides further evidence for the role of red light in mediating resistance (Pham et al., 2018), as a quadruple mutant of PIF genes, which are involved in red light signaling, is not observed in the *Arabidopsis* blue light signaling mutant *blus1* (Fig. 6a). *Arabidopsis pifq* mutant displayed less SA potentiation upon pathogen inoculation, which could be the reason behind the induction of water-soaking in this mutant. Further research will be required to dissect how red-light circuits are integrated into immune responses.

In summary, our study better defines a physiological mechanism for disease resistance and adds to the growing trend demonstrating that plant defenses involve starving pathogens for water and nutrients (Xin et al., 2016; Yamada et al., 2016). We provide a proof of concept showing that altering light conditions for relatively short periods of time can limit pathogen damage (Fig. 5) and suggest that this could have broad applications. While controlling photoperiods to prevent disease outbreaks is not as applicable for field production, this simple method could be applicable to controlled environment food production, which is growing quickly and is tightly linked to urban food sovereignty. At the same time, our current and previous work (Hu et al., 2022; Peng et al., 2019; Roussin-Léveillée et al., 2022) indicate that any treatment that leads to increased stomatal opening and transpiration leads to decreases in pathogen virulence. As such, strategies to prevent pathogen-induction of water-soaking could lead to broad-spectrum disease resistance.

## Materials and Methods

### Plant material

*Arabidopsis thaliana* plants were grown in Promix™ soil (PremierTech) in growth chambers either with 12 hours light/dark, 24 hours light or 24 hours dark photoperiod, with relative humidity of approximately 60% at 21°C. Light intensity was measured at 180 μmoles/m^2^/s in all growth chambers when lights were on. Four- to five-week-old *Arabidopsis* plants were used for all experiments described.

### Bacterial disease assays

*Pseudomonas syringae* pv. *tomato* DC3000 WT and mutant strains were cultured overnight at 28°C in Luria-Bertani (LB) media containing 50 mg/L of rifampicin. On the day of the infection, fresh LB media was inoculated with 0.5 μl of the overnight culture and bacteria were collected when OD_600_ reached between 0.8-1. Bacteria were centrifuged for 10 minutes at 4000g and the pellet resuspended in MgCl_2_ 10 mM. Bacterial density was adjusted to 0.2 (1 × 10^8^ CFU/ml) prior to further dilutions. Bacterial infections were carried between 14:00-15:00, or a zeitgeber time of 6:00-7:00.

All inoculations of *Arabidopsi*s leaves were performed by syringe-infiltration. Infiltrated plants were all kept under ambient humidity levels for 1-2 hours to allow water to evaporate, then domed with a plastic unit to maintain high humidity (>95% RH), unless stated otherwise.

Bacterial growth *in planta* was monitored by harvesting infected *Arabidopsis* leaves, surface sterilizing in 80% ethanol and rinsing in sterile water twice. Leaf disks were taken from three leaves from the same plant (one per leaf; total of three leaf disks) using a cork borer (0.6 mm in diameter) and ground in sterile 10 mM MgCl_2_. Three biological replicates were performed for each experiment. Colony-forming units (CFU) were determined by making serial dilutions (10^0^-10^−6^) and plating on LB plates containing 50 mg/L of rifampicin. Each dilution was plated in three technical replicates. Experiments were repeated at least three times.

### ROS quantification

Leaf disks from 4-week-old *Arabidopsis* plants were collected using a 4 mm diameter biopsy punch and placed into white 96-well plates (Corning) containing 100 μl of distilled water for 16 hours (overnight). Prior to ROS quantification, the water was removed and replaced with ROS assay solution (100 μM Luminol [Millipore-Sigma], 20 μg/mL horseradish peroxidase [Millipore-Sigma] with or without immune elicitors). Light emission was measured using a TECAN Spark® plate reader.

### Stomatal aperture assays

Leaves were cut at the base of the petiole and immediately immersed in the stomatal fixation solution (formaldehyde 4% and rhodamine 6G 0.5 μM) for 1 minute to stop stomatal movement. A quarter of each leaf was cut with a razor blade and stomata were observed by epifluorescence microscopy. Stomatal apertures were measured using OMERO software and the degree of the stomatal opening was measured as a ratio of stomatal width to length. Between 150 to 250 stomata were measured for each data point. Data collection and analysis was performed by using double-blinded standards to avoid bias.

### Chemical treatments

To assess plant immune responses, *Arabidopsis* leaves were treated with flg22 (1 μM; BioBasic Inc.) by syringe-infiltration. For BTH protection assay, *Arabidopsis* leaves were syringe-infiltrated with BTH (50 μM; Millipore-Sigma).

### RNA extraction, reverse transcription, and real-time quantitative PCR

RNA was extracted from frozen and ground leaf tissue using QIAZOL (QIAGEN) reagents, followed by on-column DNase treatment (QIAGEN), according to the manufacturer’s protocol. RNA purity was assessed with a spectrophotometer and quality by gel electrophoresis. cDNA was generated by using MMuLV-RT (Service de purification des protéines – Université de Sherbrooke).

Quantitative real-time PCR was performed with a Bio-Rad CFX96 machine. Each reaction contained 1X SYBR mix (Service de purification des protéines – Université de Sherbrooke), specific primers and a 1:20 dilution of 500 ng of cDNA stock. Amplification cycle protocols were as follow: 2 min at 95°C; 40 cycles of 6 seconds at 95°C and 30 seconds at 60°C. Melting curves were verified at the end of 40 cycles for confirmation of primer specificities. All reactions were repeated in three technical and biological replicates. Average Cq values were normalized by ΔΔCT formula against the indicated reference gene. Oligos used in this study can be found in Table 1.

### Apoplast extraction

Three fully expanded leaves per plant from three plants were excised and apoplast extracted as previously described (Gentzel et al., 2019), with some modifications. Briefly, the initial weight of freshly excised leaves was measured before being infiltrated with distilled water and weighed again once leaves were fully saturated with water. Leaves were centrifuged at 4000 rcf in 2 ml microcentrifugation tubes containing glass beads at the bottom to prevent tissue collapse during the centrifugation. Leaves were once again weighed post centrifugation and apoplast hydration determined as previously described (Gentzel et al., 2019).

### Salicylic acid extraction and quantification

Fully expanded four-week-old *Arabidopsis* leaves were harvested and weighed for fresh weight calculation and immediately flash-freeze in liquid nitrogen. Tissues were ground with a plastic pestle and phytohormones were extracted overnight using 0.5-1 ml of ice-cold extraction buffer (methanol: water [80:20 v/v], 0.1% formic acid, 0.1 g/L butylated hydroxytoluene and 100 nM ABA-d6 as an internal standard). Extracts were filtered using centrifugal filter units.

Filtered extracts were quantified using an Acquity Ultra Performance Liquid Chromatography system (Waters Corporation, Milford, MA) as described previously (Roussin-Léveillée et al., 2022). SA was quantified based on a standard curve to calculate sample concentration (nM), which was converted to ng using the molecular weight of each specific compound and the extraction volume used. All data was normalized to initial fresh weight in grams.

### Statistical analysis

Statistical analyses were performed in GraphPad Prism 8.0 for all RT-qPCR experiments, photon counts, and SA quantification. For all other experiments (bacterial counts and stomatal assays), the bioinformatic software R was used. *P* values greater than 0.05 were considered non-significant. Sample sizes, statistical tests used, and *P* values are stated in figure legends. All statistical test assumptions such as normality and homoskedasticity were tested. When not respected, non-parametric equivalent tests were performed. In multiple comparison tests, Bonferroni corrections were used.

## Acknowledgments

This study was supported by two National Sciences and Engineering Research Council of Canada (NSERC) Discovery Grants respectively to P.M. and I.L.-L., as well as by a Fonds de Recherche du Québec – Nature et Technologies (FRQ-NT) Team Grant and NOVA-NSERC Alliance Grant to P.M. (PI) and I.L.-L. (co-PI). I.L.-L. is also supported by a Canada Research Chair T2. G.L. was supported by an Excellence Scholarship from the Faculty of Science of Université de Sherbrooke and by a Canada Graduate Scholarship (NSERC-MSc). C.R.-L. was supported by a VoiceAge excellence fellowship and an FRQ-NT PhD scholarship.

## Author contributions

C.R.-L. and P.M. conceived the study. G.L. and C.R.-L. performed all experiments, data analysis and the creation of figures. S.B. and E.F. contributed to some experiments and data collection. I.L.-L. contributed to the statistical analysis. G.L. and C.R.-L. wrote the manuscript with support from P.M. and I.L.-L. All authors revised and agreed with the content written in this manuscript.

