## Supplemental Figures for "Light prevents pathogen-induced aqueous microenvironments via potentiation of salicylic acid signaling"

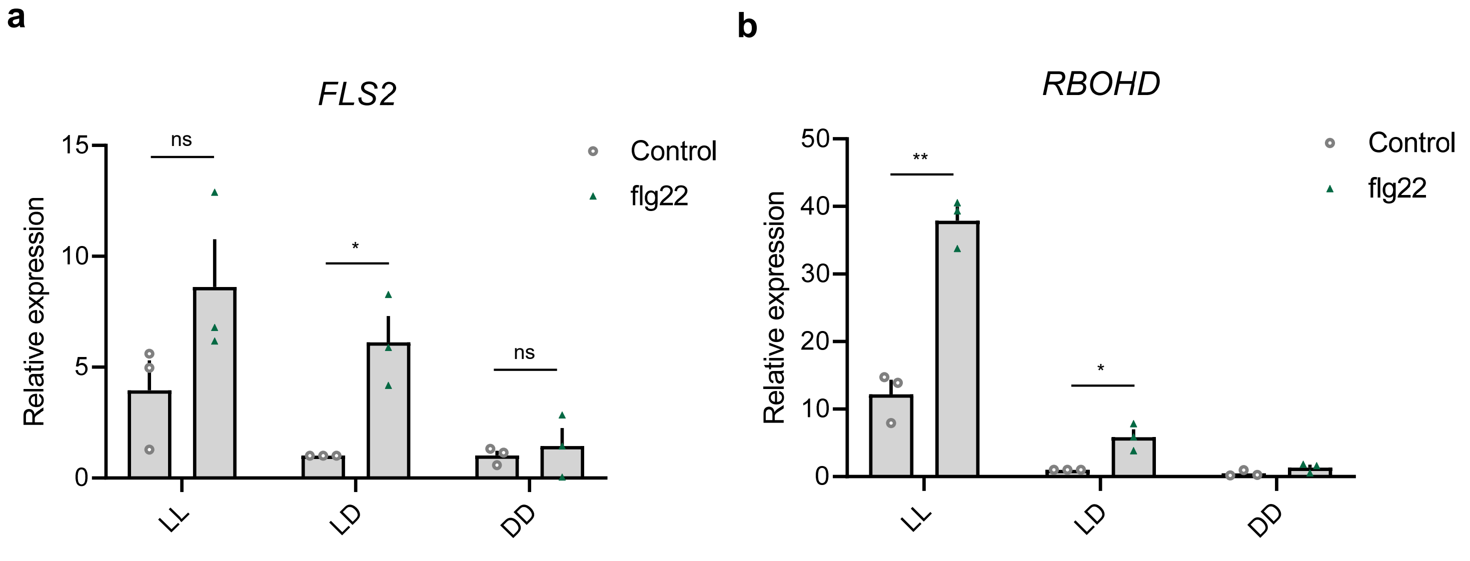


**Figure S1: Expression of plasma-membrane localized PTI associated genes under different light regimes**

**a-b,** Expression levels, as determined by qRT-PCR, of *FLS2* (**a**) and *RBOHD* (**b**) in WT *Arabidopsis* plants that were subjugated to 24 hours of the indicated light cycles prior to being infiltrated with flg22 (1 μM). Samples were harvested 6 hours post flg22 challenge. Asterisks indicate statistically significant differences compared to control. ns = non-significant, $*p < 0.05$, $*p < 0.005$ Student's T-test

**a**

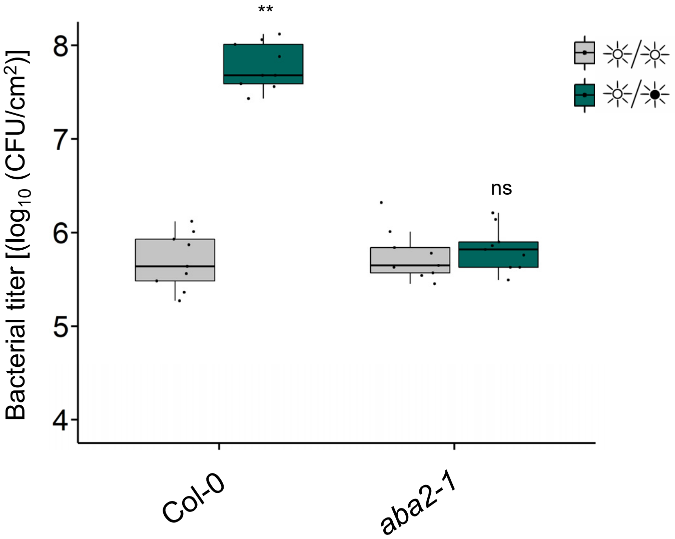


**Figure S2: Impact of ABA on dark-induced virulence**

**a**, Bacterial titers from WT *Arabidopsis* and *aba2-1* mutant plants syringe-infiltrated with *Pst* DC3000 ($1x10^5$ CFU/ml) at 3 dpi, under the indicated light regimes. Asterisks indicate statistically significant differences compared to control. ns = non-significant, Student's t test (*aba2-1*) or Wilcoxon-Mann Witney test (Col-0).

**a**

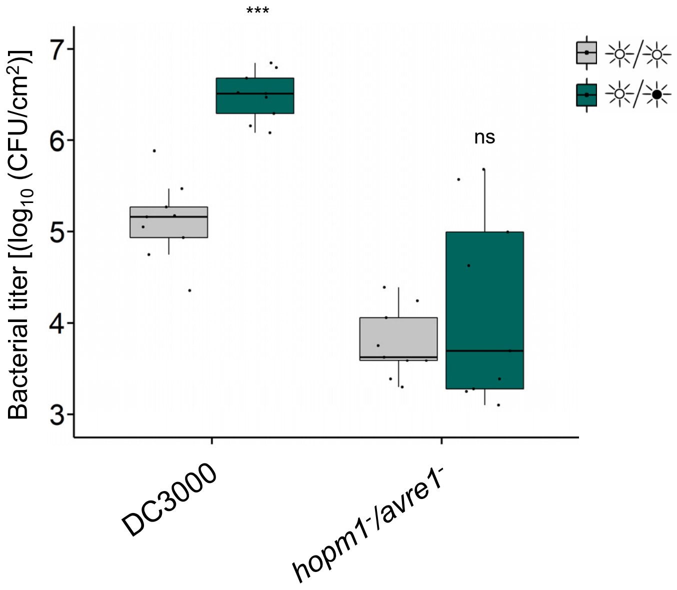


**Figure S3: HopM1 and AvrE1 contribution to disease progression requires darkness**

**a**, Bacterial titers from WT *Arabidopsis* plants syringe-infiltrated with *Pst* DC3000 (1x$10^5$ CFU/ml) and the water-soaking effector mutant *Pst hopm1⁻/avre1⁻* at 3 dpi under the indicated light regimes. ns = non-significant, *** $p < 0.0001$ , Student's t test (DC3000) or Wilcoxon-Mann Witney test (*hopm1⁻/avre1⁻* ).

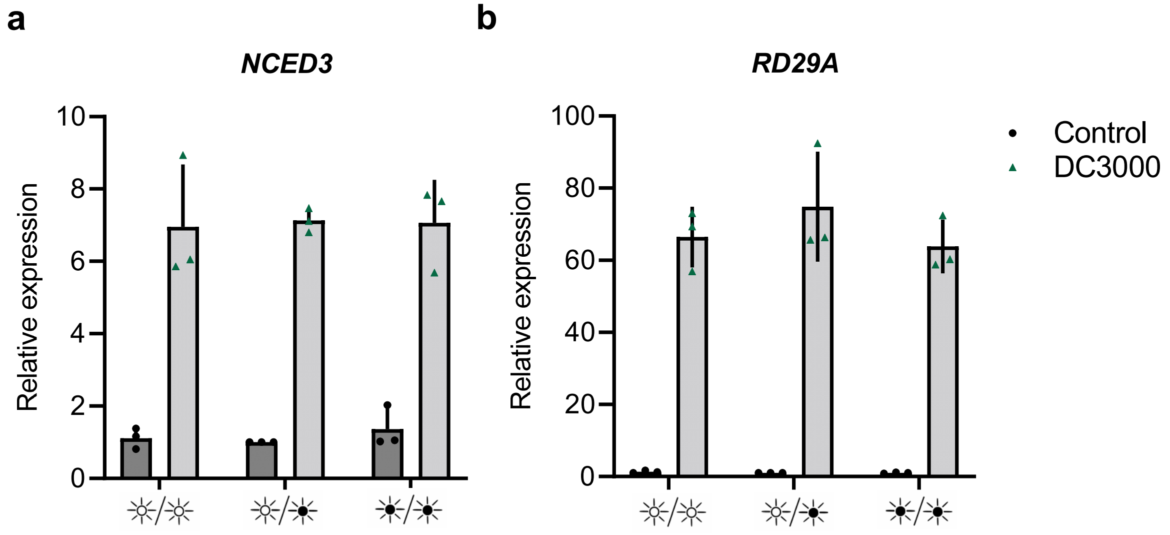


**Figure S4: ABA biosynthesis and signaling is not affected by light regimes upon infection**

**a-b**, Expression levels, as determined by qRT-PCR, of ABA biosynthesis (*NCED3*) and signaling (*RD29A*) marker genes in WT *Arabidopsis* plants, kept under the indicated light regimes, infected with *Pst* DC000 ($1x10^8$ CFU/ml) or $MgCl_2$10 mM (control) at 24 hpi.

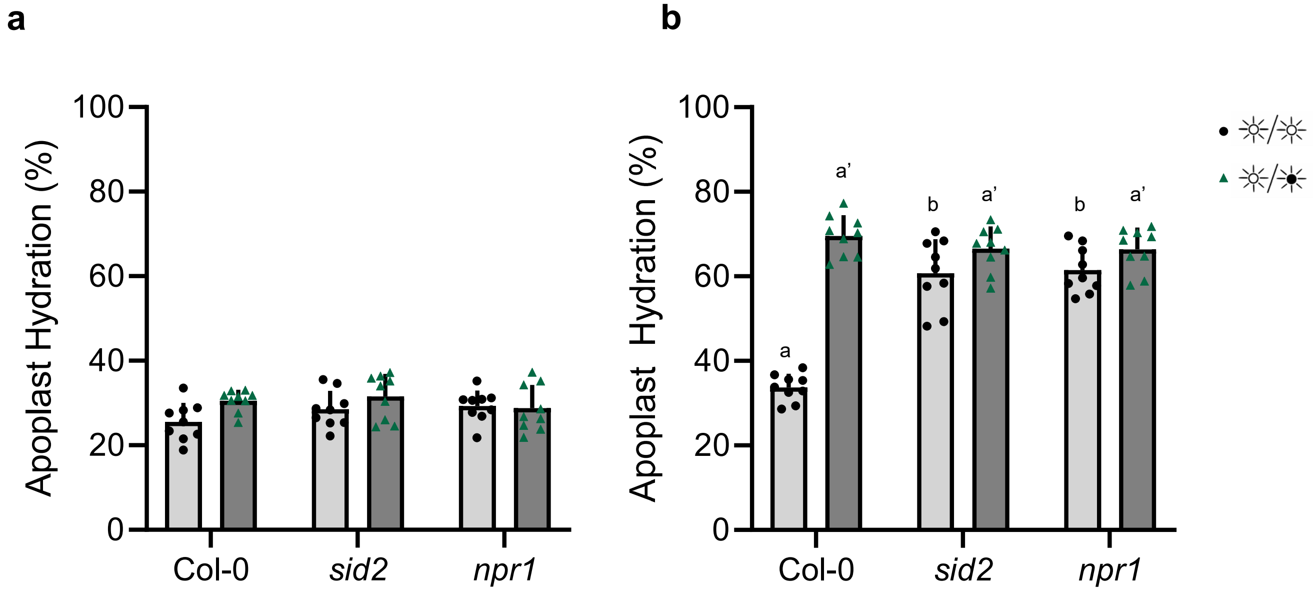


**Figure S5: Apoplast hydration in SA biosynthesis and signaling mutant plants under different light regimes**

**a-b**, Apoplast hydration levels at 24 hpi measured from four-week-old *Arabidopsis* WT (Col-0), *sid2* and *npr1* mutant plants infiltrated with 10 mM $MgCl_2$ (**a**) or with $1x10^8$ CFU/ml of *Pst* DC3000 (**b**). Plants were put under constant light for the whole 24 hours period or kept under a 12 hour light/dark regime. Different letters indicates statistically significant differences, $p < 0.05$, ANOVA followed by Tukey's range test.

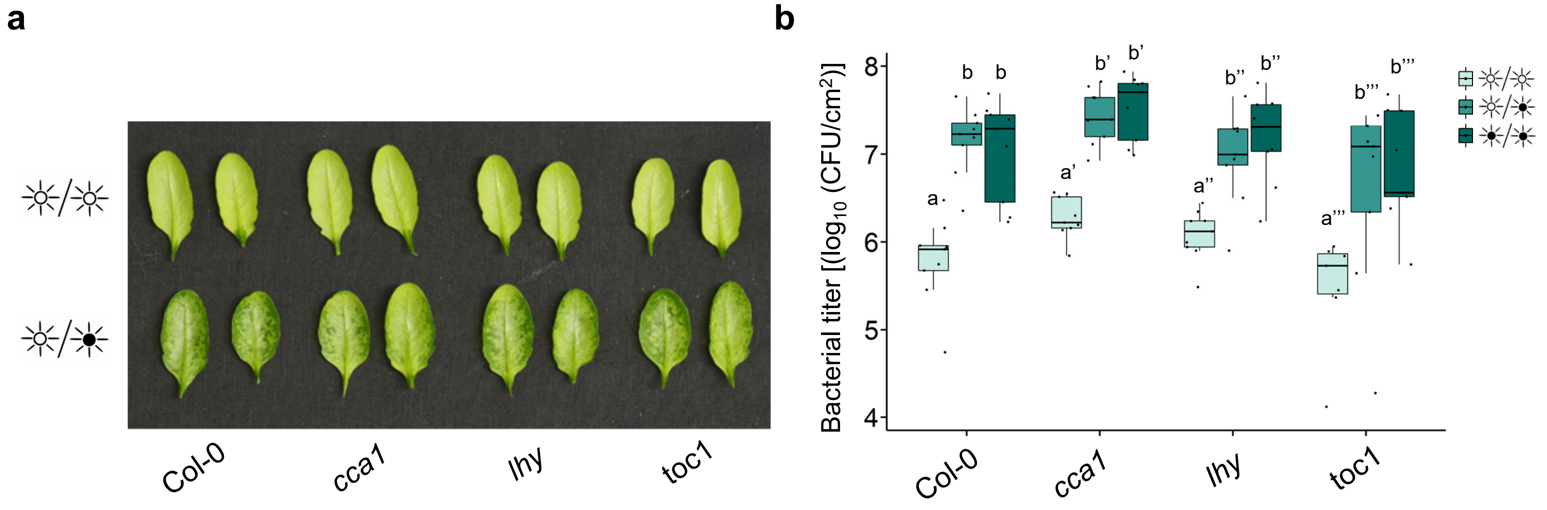


**Figure S6: Circadian clock mutants are not affected in water-soaking lesions under constant light**

**a,** Water-soaking phenotypes of *Arabidopsis* WT (Col-0), *cca1*, *lhy* and *toc1* mutant plants syringe-infiltrated with *Pst* 3000 ($1 \times 10^8$ CFU/ml) under the indicated light settings. Photos were taken at 24 hpi. **b**, Bacterial titers from *Arabidopsis* WT (Col-0), *cca1*, *lhy* and *toc1* mutant plants syringe-infiltrated with *Pst* 3000 ($1 \times 10^5$ CFU/ml) under the indicated light settings at 3 dpi. Different letters indicate statistically significant differences, $p < 0.05$, ANOVA (Col-0, *lhy*) or Kruskal-Wallis test (*cca1*, *toc1*).
